# Genome-wide patterns of gene expression in a wild primate indicate species-specific mechanisms associated with tolerance to natural simian immunodeficiency virus infection

**DOI:** 10.1101/395152

**Authors:** Noah D. Simons, Geeta N. Eick, Maria J. Ruiz-Lopez, David Hyeroba, Patrick A. Omeja, Geoffrey Weny, Colin A. Chapman, Tony L. Goldberg, HaoQiang Zheng, Anupama Shankar, William M. Switzer, Simon D.W. Frost, James H. Jones, Kirstin N. Sterner, Nelson Ting

## Abstract

Over 40 species of nonhuman primates host simian immunodeficiency viruses (SIVs). In natural hosts, infection is generally assumed to be nonpathogenic due to a long coevolutionary history between host and virus, although pathogenicity is difficult to study in wild nonhuman primates. We used whole-blood RNA-seq and SIV prevalence from 29 wild Ugandan red colobus (*Piliocolobus tephrosceles*) to assess the effects of SIV infection on host gene expression in wild, naturally SIV-infected primates. We found no evidence for chronic immune activation in infected individuals, suggesting that SIV is not immunocompromising in this species, in contrast to HIV in humans. Notably, an immunosuppressive gene, *CD101*, was upregulated in infected individuals. This gene has not been previously described in the context of nonpathogenic SIV infection. This expands the known variation associated with SIV infection in natural hosts, and may suggest a novel mechanism for tolerance of SIV infection in the Ugandan red colobus.

## INTRODUCTION

The effects of infectious pathogens on host physiology are often influenced by coevolution between host and pathogen. This understanding can provide insight into aspects of adverse health outcomes in humans, particularly in cases where host-pathogen relationships can be analyzed in a comparative perspective (Enard et al. 2010, 2016). A prominent example is the human immunodeficiency viruses (HIVs), the causative agents of acquired immunodeficiency syndrome (AIDS), which have close relatives in nonhuman primates (Gao et al., 1999; Hirsch et al., 1989). The study of coevolution between African primates and simian immunodeficiency virus (SIV) has provided much insight into our understanding of HIV (Palesh et al., 2018). In contrast to HIV, SIV infection in natural hosts is generally considered nonpathogenic, reflecting a long history of coevolution favoring benignness in the virus (Chahroudi et al., 2012; Jasinska et al., 2013). Over 40 species of nonhuman primates are naturally infected with endemic strains of SIV, including African apes and African members of the Old World monkey subfamilies Cercopithecinae and Colobinae (VandeWoude & Apetrei, 2006; Worobey et al., 2010). However, studies on the pathogenicity of SIV infection in natural hosts have been limited to chimpanzees and three species within the Cercopithecinae (African green monkey (AGM), sooty mangabey (SM), and mandrill (MND; Apetrei et al., 2011; Bosinger et al., 2009; Greenwood et al., 2014; Jacquelin et al., 2009; Keele et al., 2009; Ma et al., 2013). Recently, studies in SIVcpz-infected chimpanzees (Keele et al., 2009) and SIVmnd-1-infected MNDs (Greenwood et al., 2014) have demonstrated variation in the pathogenic effects of SIV infection in natural hosts, but to our knowledge no study has assessed the relationship between SIV infection and host gene expression patterns in a wild population of natural SIV hosts. It has become increasingly clear that the effects of SIV infection vary across natural hosts, and the assumption of nonpathogenic SIV infection in natural hosts warrants further testing in a wider breadth of taxa, particularly in understudied groups such as the Colobinae (Bosinger et al., 2013; Palesh et al., 2018).

### Nonpathogenic SIV infection

SIV/HIV infection can be broadly divided in to two phases: acute (weeks) and chronic (years). The immune response during acute SIV infection in natural hosts parallels that of SIV/HIV-infected non-natural hosts (macaques and humans) in several ways. During the acute phase, both pathogenic and nonpathogenic infection are characterized by a rapid induction of Type I interferon response (IFN; signaling proteins produced in response to viral infections that initiate a signaling cascade of downstream antiviral effector proteins; Sadler & Williams, 2008), upregulation of IFN-stimulated gene (ISG) expression, high viral loads, and a decrease in peripheral CD4+ T cell count (reviewed by Chahroudi et al. 2012). Nonpathogenic SIV infection is distinguished from pathogenic infection during the transition from acute to chronic phase, approximately 21-40 days after infection (Mir et al., 2011). During this period, one of the hallmark features of nonpathogenic SIV infection is the rapid attenuation of ISG expression to baseline levels in the chronic phase (Bosinger et al. 2009; Jacquelin et al. 2009; Lederer et al., 2009). Additionally, natural hosts maintain high viral loads, but recover CD4+ T cell homeostasis, though variation in the effects of SIV infection on CD4+ T cell homeostasis have been observed in natural hosts (Keele et al. 2009; Greenwood et al., 2014). While the upregulation of ISGs persists in pathogenic SIV/HIV infection in macaques and humans (Paiardini & Müller-Trutwin, 2013), rapid control of ISG expression occurs in natural hosts (AGMs and SMs) within 4-8 weeks post-infection (Bosinger et al. 2009; Jacquelin et al. 2009). The gene expression patterns underlying this downregulation of ISGs have only been investigated in experimentally infected captive populations of two species, AGMs and SMs. Research on AGMs and SMs suggests that natural hosts actively downregulate the innate and adaptive immune response following infection (Chahroudi et al. 2012), and studies have proposed negative regulatory mechanisms by which the acute immune response is attenuated through the upregulation of a small number of immunosuppressive genes, including *IDO, LAG3, IL10*, and *LGALS3* in AGMs (Jacquelin et al. 2009) and *ADAR* in SMs (Bosinger et al. 2009). A recent study by Svardal and colleagues (2017) also found a high degree of overlap between genes under selection in the AGM genome and genes differentially expressed in AGMs, but not macaques, in response to SIV infection (using microarray data from Jacquelin et al. 2009).

### SIVkrc infection in the Ugandan red colobus

Natural populations of Ugandan red colobus (URC; *Piliocolobus tephrosceles*; Old World monkey subfamily Colobinae) have emerged as an important wild primate species for understanding the role of host biology in viral disease (Bailey et al., 2014, 2016a, 2016b; Goldberg et al., 2008, 2009; Ladner et al., 2016; Lauck et al., 2011, 2013; Sibley et al., 2014). SIV diversity in red colobus was first described in western red colobus (*Piliocolobus badius*) in Taï forest in Côte d’ Ivoire and established the red colobus as a potential reservoir for viral transmission to humans (Locatelli et al., 2008a). Full SIV genome characterization from a different western red colobus (*P. temminckii*) from the Abuko Nature Reserve, The Gambia, showed that SIV lineages in red colobus were species-specific (Locatelli et al., 2008b). Goldberg and colleagues (2009) showed that URC in Kibale National Park, Uganda are natural hosts for a novel SIV lineage designated SIVkrc.

Genomic studies of host responses to SIV infection in natural populations have been limited by several factors, particularly the difficulty of obtaining and preserving high quality biomaterials from animals that exist in remote locations and are often endangered. Transcriptomic approaches to understanding the effects of SIV infection on host gene expression patterns have therefore been limited to studies of captive populations of AGMs and SMs, in which the effects of experimental infection have been assessed with the use of microarrays (Bosinger et al. 2009; Jacquelin et al. 2009). Those studies provide valuable hypotheses, which can now be tested in wild populations of naturally infected hosts, to examine the effects of SIV infection in natural hosts within the ecological context of host-virus coevolution.

In this study we used gene expression profiles from SIVkrc-infected and SIVkrc-uninfected URC from Kibale National Park, Uganda to address three questions. (1) Do ISG expression patterns differ between naturally SIVkrc-infected and uninfected URC? Both AGMs and SMs mount an acute ISG response that is attenuated in the chronic phase (Bosinger et al. 2009; Jacquelin et al. 2009). Given that signs of immunodeficiency are difficult to observe in URC, and our study subjects are presumed to be in the chronic phase of infection, we hypothesize that levels of ISG expression will be attenuated to baseline levels similar to uninfected URC. (2) Is the expression of immunosuppressive genes elevated in SIVkrc-infected URC relative to uninfected URC? Upregulation of immunosuppressive genes upon infection and sustained through the chronic phase has been observed in AGMs and SMs (Bosinger et al. 2009; Jacquelin et al. 2009). We hypothesize that SIVkrc-infected URC will show upregulation of immunosuppressive genes compared to uninfected URC. (3) Do SIVkrc-infected URC show CD4+ T cell depletion compared to uninfected URC? The effect of SIV infection on T cell subset populations varies across natural hosts, as AGMs and SMs maintain CD4+ T cell levels, while SIVmnd-1-infected MNDs and SIVcpz-infected chimpanzees show a loss of memory CD4+ T cells (Chahroudi et al. 2012; Greenwood et al. 2014; Keele et al. 2009), which provide immunological memory as part of the adaptive immune response (Harrington et al., 2008). Given the different SIV infection patterns in AGMs, SMs, MNDs, and chimpanzees, we test the null hypothesis that SIVkrc-infected URC will maintain CD4+ T cell levels comparable to uninfected URC.

This study represents the first high-resolution transcriptomic analysis of the relationship between SIV infection and host gene expression in a wild population of naturally infected SIV hosts. It is also the first study on the host gene expression patterns in relation to SIV infection in a colobine, and adds considerably to our current knowledge about variation in host patterns of gene expression across natural SIV hosts.

## RESULTS

### Sex differences in SIVkrc prevalence but not viral load

To test the effects of SIVkrc infection on patterns of gene expression we generated SIVkrc prevalence and viral load data from 29 individuals using two measures: 1) SIV antibody detection using HIV-2 western blot assays, and 2) presence of SIV RNA envelope (*env*) and LTR sequences using two SIVkrc-specific RT-qPCR assays. Both measures of SIV infection (serology and RT-qPCR) were 100% concordant. SIVkrc infection prevalence in our study group was 41.4% (Wilson’s 95% confidence interval (CI) = 25.5%-59.3%; Wilson, 1927), with females having a substantially lower prevalence than males (F = 25% (CI = 7.1%-59.1%), M = 52.6% (CI = 31.7%-72.7%); Supplemental Table S1). Our observation of lower prevalence in females than males parallels that reported by Goldberg and colleagues (2009), and we observed an increase in overall prevalence of 18.8% since 2009.

Viral load (reported as log_10_ genome copies per mL of plasma) estimates were similar between the *env* and LTR RT-qPCR assays (Mann Whitney *U* = 94, *Z* = −1.24, *p* = 0.22; Figure 1). There were no significant differences between males and females for either *env* (*t*(10)= 1.06, *p* = 0.31) or LTR (*t*(10) = 0.66, *p* = 0.26), and viral load for both measures were within the range of those reported using RT-qPCR of the viral *int* protein for three species of naturally SIV-infected AGMs (Bailey et al. 2016a; Ma et al., 2014; Figure 1).

**Figure 1.**
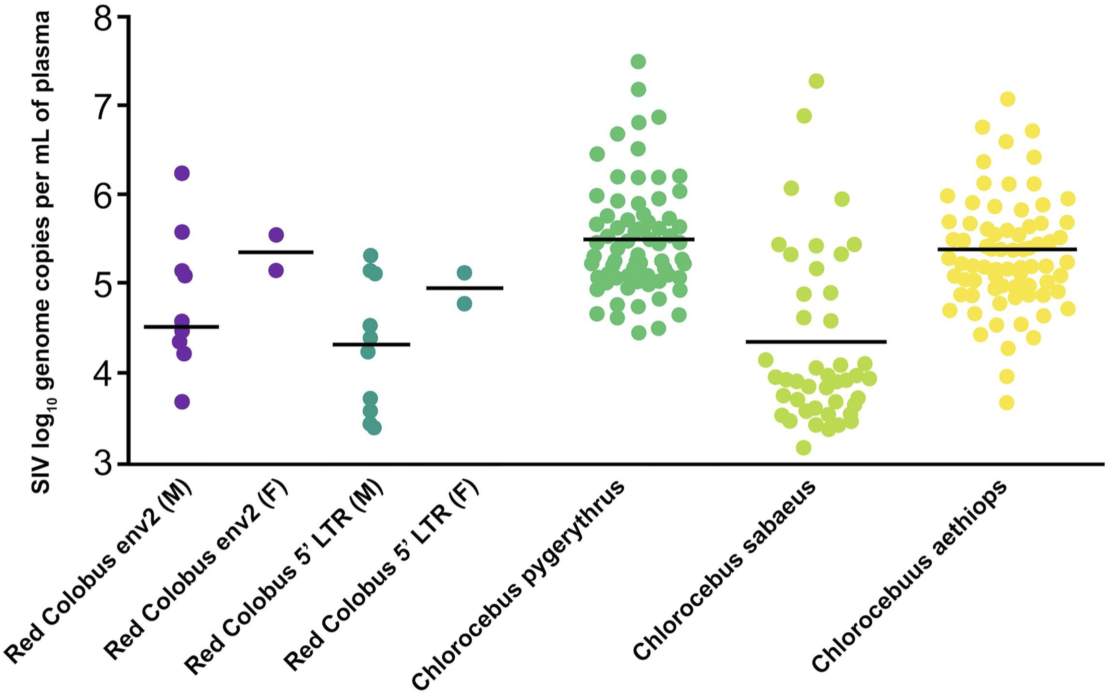
Viral load (reported as log_10_ genome copies per mL of plasma) generated with RT-qPCR of *env* and LTR for red colobus. Viral load data for *C. pygerythrus* from (Bailey et al. 2016a), *C. sabaeus* and *C. aethiops* from (Ma et al. 2014).

### Sequence data, mapping, and quantification

To identify gene expression differences between SIVkrc-infected and uninfected URC, we quantified genome-wide gene expression from the same 29 URC individuals using RNA-seq. We generated 984.7 million 150 base-pair reads (average ∼33.95M reads per individual). We retained 871.2 million reads (average ∼30.04M reads per individual) after filtering. A high proportion of reads were successfully mapped to the most recent version of the annotated URC reference genome (GCF_002776525.1), with an average of 23.7M (78.3%, *SD* = 7.6) reads mapping uniquely, and average of 2.1M (6.6%, *SD* = 1.6) reads mapping to multiple locations. Of uniquely mapped reads, an average of 16.2M (67.8%, *SD* = 0.08) reads mapped to gene features in the red colobus annotation. Total mapped (combined unique and multi-mapped) reads averaged 25.7M (85.8%, *SD* = 8.5). The number of annotated genes expressed in whole blood of the URC (N = 11,452) was consistent with that found in humans (N = 12,758; Göring et al., 2007), baboons (N = 10,409; Tung et al., 2015), and wolves (N = 13,558; Charruau et al., 2016).

### Individual gene expression patterns do not cluster according to infection status

We used two sample-based clustering methods to determine if individual gene expression patterns clustered according to infection status. After VST transformation, a total of 17,583 genes (49.1% of original 35,827) were retained for clustering and visualization. Both hierarchical clustering based on sample distances (Figure 2), and PCA (Figure 3) indicated that individuals did not cluster according to infection status (infected vs. uninfected) or sex (M vs. F). Pathway analysis of loadings from PC1 found strong enrichment of pathways related to RNA processing (*Q* = 1×10e-292), cellular protein (*Q* = 1×10e-284), macromolecule localization (*Q* = 4×10e-287), and organelle organization (*Q* = 1×10e-293; Supplemental Table S2). Pathway analysis of loadings from PC2 found that pathways related to ribosome biogenesis (*Q* = 1×10e-56) and ncRNA processing (*Q* = 1×10e-64) were enriched.

**Figure 2.**
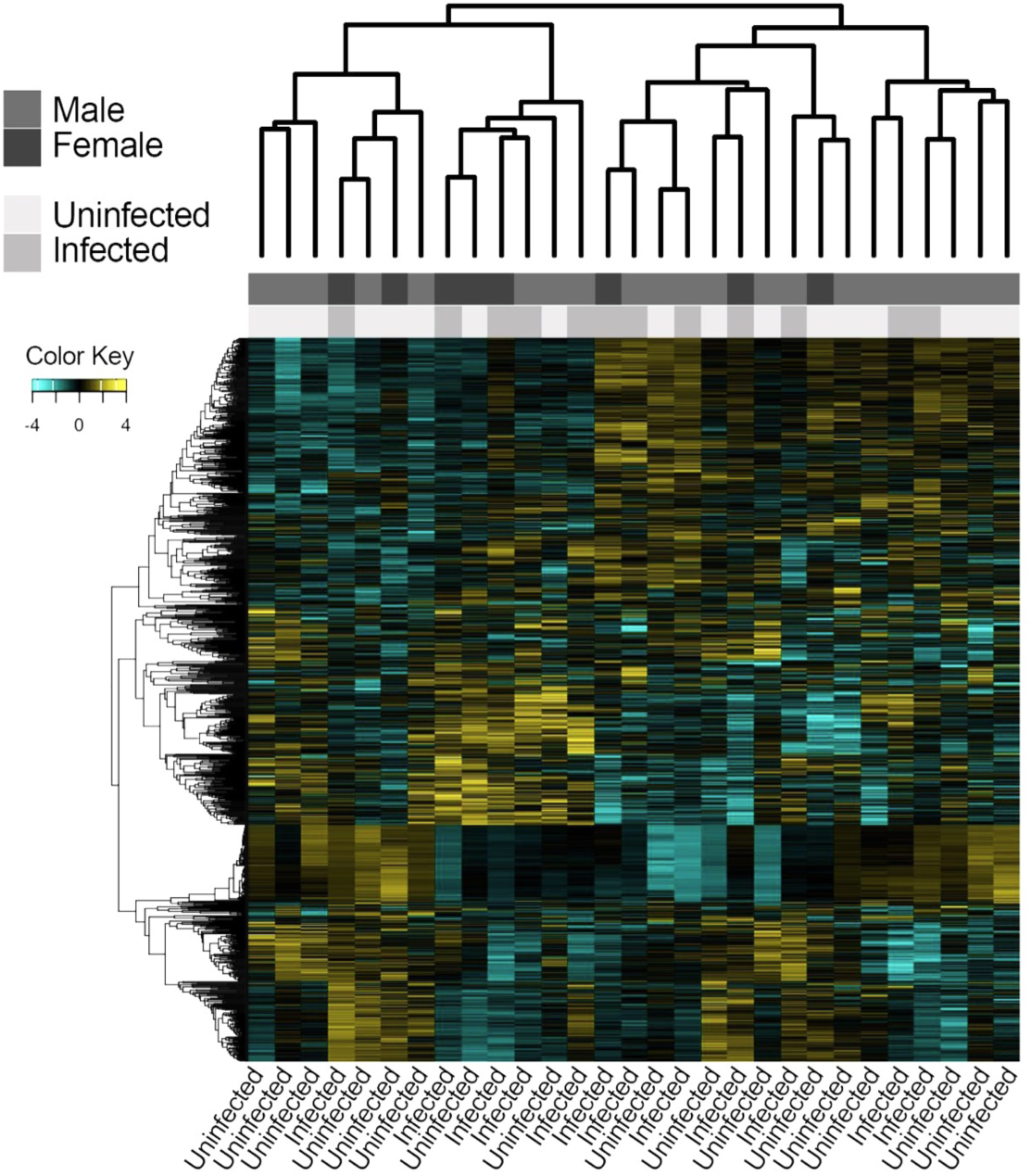
Hierarchical clustering showing that individuals do not cluster according to infection status or sex.

**Figure 3.**
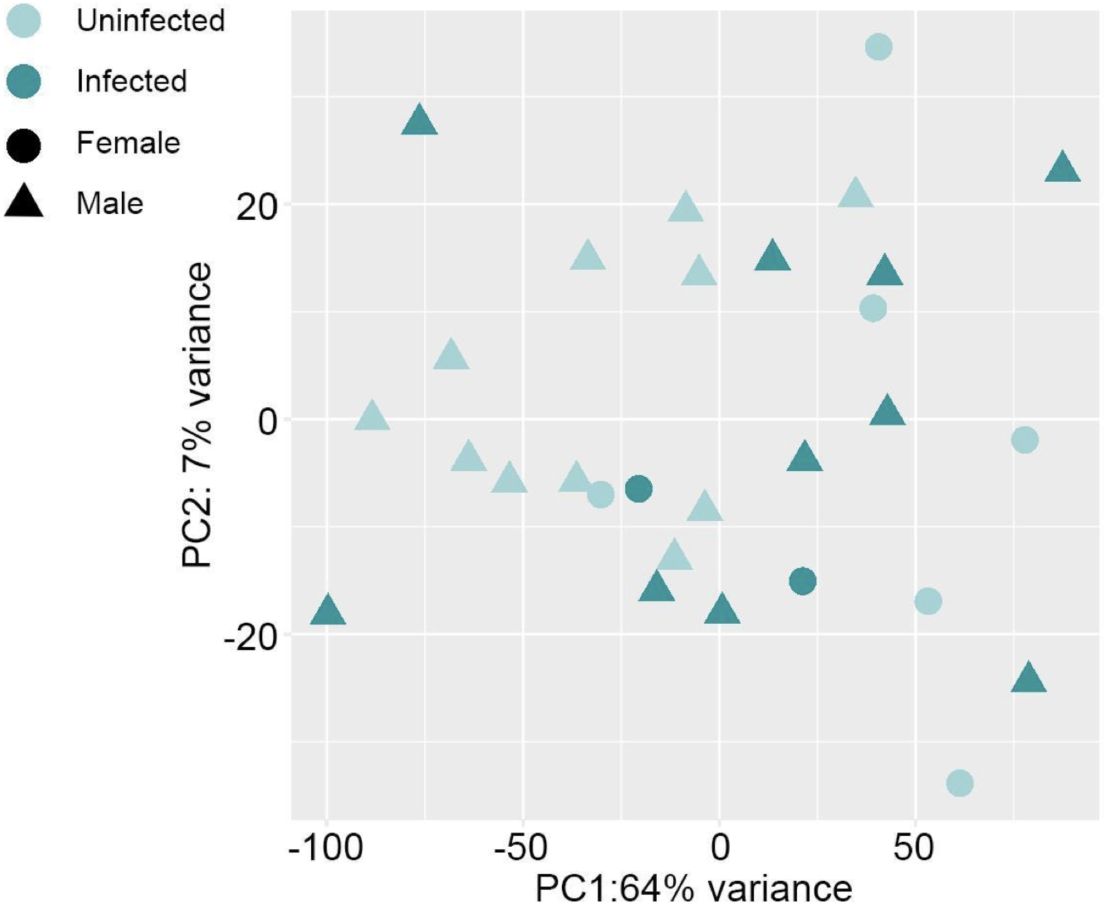
Principal component analysis (PCA) of gene expression patterns showing that neither PC1 or PC2 separate samples according to infection status or sex.

### K-means clustering of genes identifies fewer enriched immune pathways in SIVkrc-infected individuals than uninfected individuals

We used gene clustering to assess if infected individuals had more gene clusters enriched for immune-related pathways than uninfected individuals. K-means clustering identified five clusters of correlated genes in uninfected individuals (Figure 4), and four of the five (A, C, D, and E) were enriched primarily with immune related pathways (Supplemental Table S3A). Four clusters were identified (A-D) in infected individuals, of which two clusters (C and D) were enriched with immune related pathways (Supplemental Table S3B). Promoter analysis of clusters in uninfected individuals found 17 enriched transcription factor binding motifs in three clusters (B, C, and D). Enriched binding motifs consisted of four transcription factor families: C2H2 ZF, bHLH, Rel, and SMAD (Supplemental Table S4A). Promoter analysis in infected individuals found 38 enriched transcription factor binding motifs, consisting of seven transcription factor families: C2H2 ZF, E2F, Homeodomain, MBD, Myb/SANT, bHLH, Rel (Supplemental Table S4B). Three transcription factor families (C2H2 ZF, bHLH, Rel) found in uninfected individuals were also found in infected individuals, suggesting global enrichment of those three families in the blood transcriptome of URC, regardless of infection status. Four transcription factor families were unique to infected individuals (MBD, Myb/SANT, Homeodomain, and E2F), and may play a role in the downregulation of the immune response during infection, as the specific transcription factors of three of the four (MBD, Myb/SANT, and E2F) families are transcriptional repressors.

**Figure 4.**
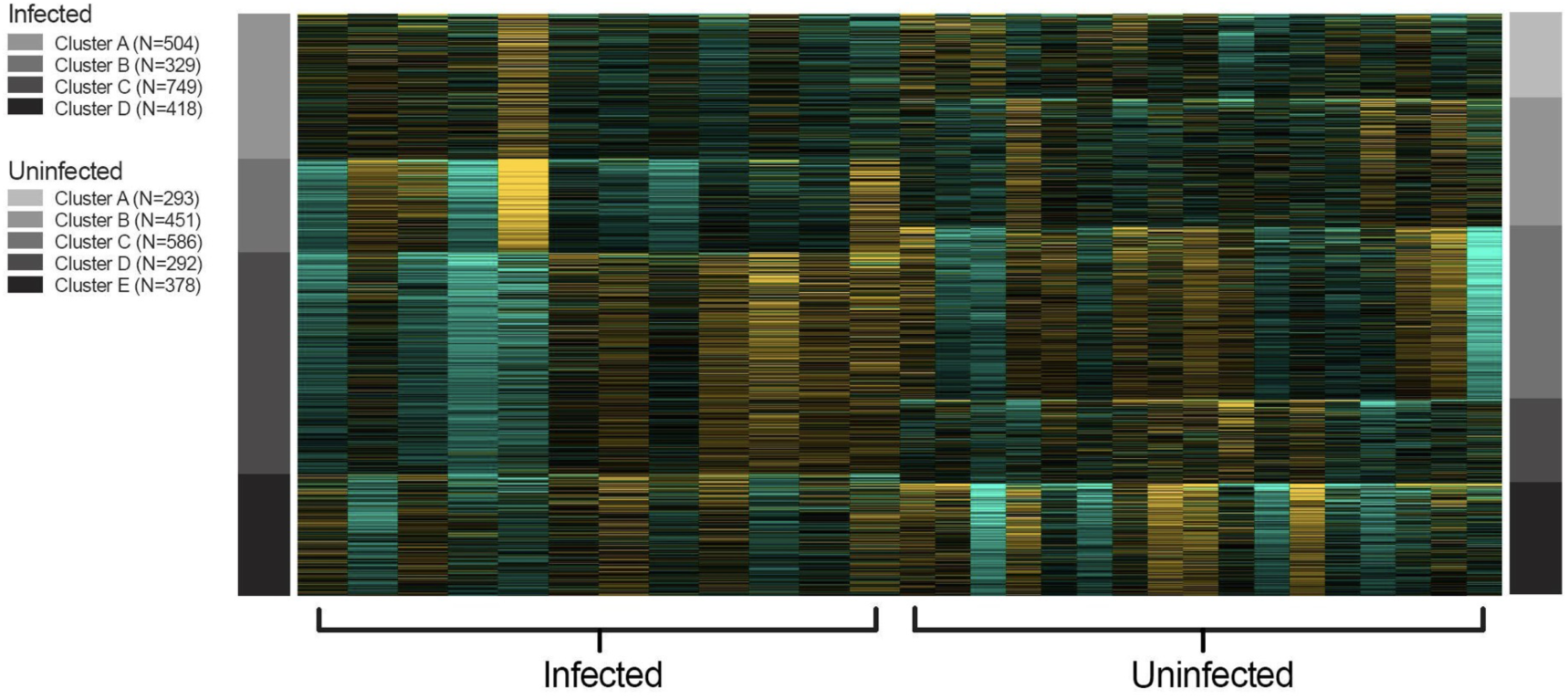
K-means clustering of top 2000 most highly variable genes. K-means identified four and five gene clusters in infected and uninfected individuals, respectively.

### Global signatures of immune activity in the blood transcriptome

We used gene co-expression networks to identify correlated networks of genes and assess if those networks were enriched for immune-related pathways in infected individuals compared to uninfected individuals. Weighted gene co-expression network analysis identified a network of 2872 genes divided into 17 co-expression modules in infected individuals, and a network of 2886 genes divided into 11 co-expression modules in uninfected individuals (Figure 5). GO (Gene Ontology) pathway enrichment analysis found several highly enriched immune-related pathways for both infected and uninfected individuals, including immune system process, cell activation, response to external stimulus, leukocyte activation, immune response, and immune effector process, likely reflecting global patterns of immune function in the blood transcriptome. Immune-related GO pathways enriched only in infected individuals were all related to cellular immunity, including cell activation involved in immune response, leukocyte activation involved in immune response, and myeloid leukocyte activation. In contrast, the only GO pathway enriched in uninfected individuals, but not in infected individuals, was inflammatory response (Supplemental Table S5).

**Figure 5.**
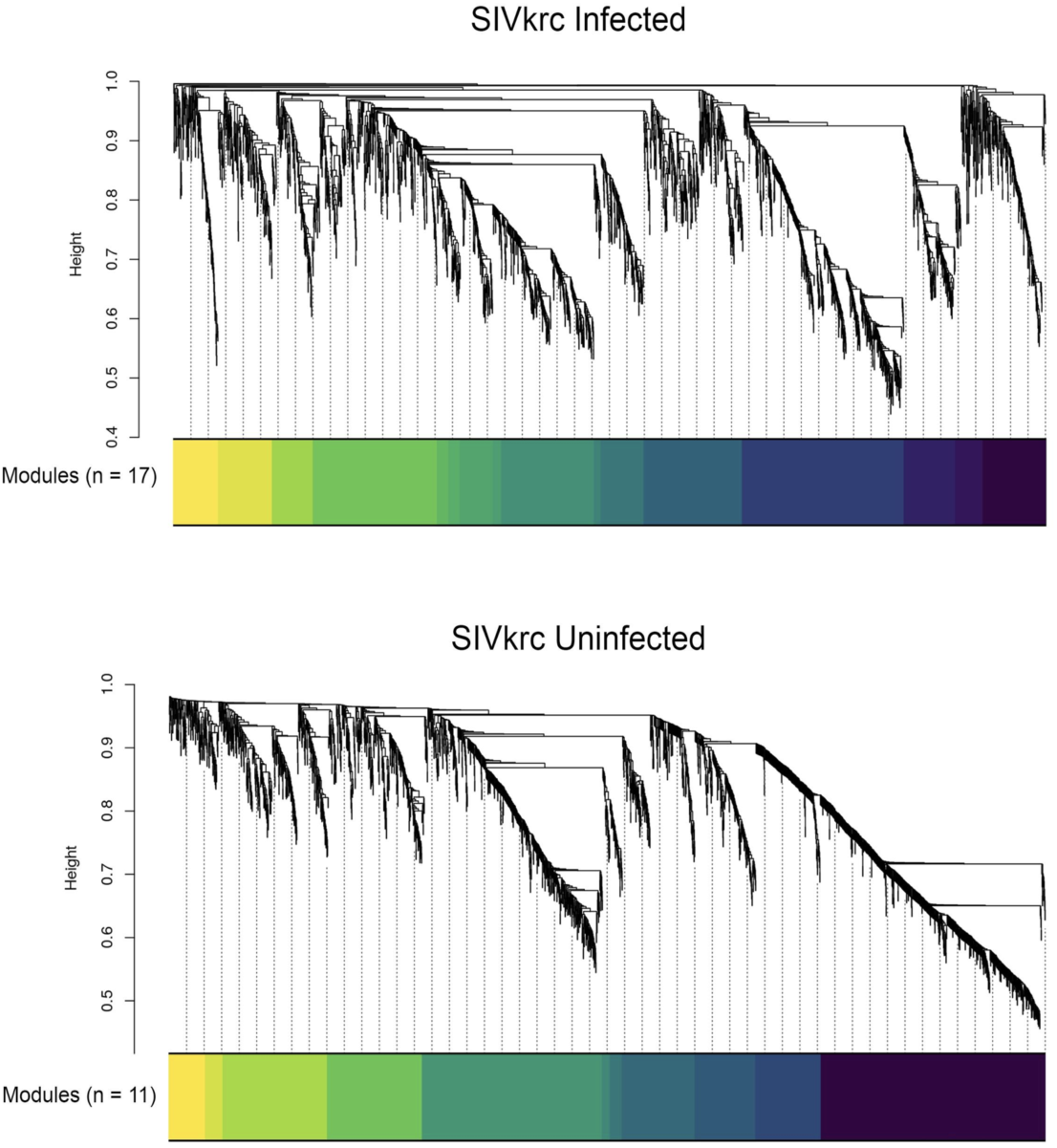
Weighted gene co-expression network analysis identified a network of 2,872 genes divided into 17 co-expression modules in infected individuals, and a network of 2,886 genes divided into 11 co-expression modules in uninfected individuals.

### Attenuated ISG and upregulated immunosuppressive gene expression in SIVkrc-infected URC

We modeled the effects of SIVkrc infection on gene expression and identified 53 differentially expressed genes between SIVkrc-infected and uninfected individuals, with eight genes downregulated and 45 genes upregulated in infected individuals (Supplemental Table S6). Genes downregulated in infected individuals included two immune genes, *SLAMF6* (ENSG00000162739; fold change = −0.39; *Q* value = 0.12) and *CD4* (ENSG00000010610; fold change = −1.58; *Q* value = 3×10e5). For both of these genes, there is evidence of virus-mediated down-modulation following infection. For example, for *SLAMF6* (a.k.a. *NTB-A*), the HIV-1 accessory protein *Vpu* down-modulates *SLAMF6* on infected T cell surfaces, thereby protecting infected cells from lysis by natural killer cells (Shah et al., 2010). Interestingly, SIVkrc lacks the *Vpu* accessory protein (Sakai et al., 2016). For *CD4*, which is the primary cell-entry receptor for both SIV/HIV, the viral accessory protein *Nef* down-modulates cell-surface expression of *CD4* through endocytosis (Pham et al., 2014). Two GO terms were enriched in downregulated genes in infected individuals: positive regulation of biological process (*Q* value = 4.65E-04) and cytosolic calcium ion concentration (*Q* value = 4.99E-03; Table 1). GO terms enriched in genes upregulated in infected individuals included broad categories related to cellular processes and biological regulation. There were no enriched GO terms related to immune response in significantly up or downregulated genes in infected individuals (Table 1; full GO results in Supplemental Table S7).

**Table 1.** GO terms enriched in up and down-regulated genes in SIVkrc infected individuals. Corrected *p* values are g:SCS corrected. See full GO results in Supplemental Table S7.

| Up |  | Down |  |
| --- | --- | --- | --- |
| Top Enriched GO Term | Corrected GO p Value | Top Enriched GO Term | Corrected GO p Value |
| cellular metabolic process | 8.09E-09 | positive regulation of biological process | 4.65E-04 |
| vesicle-mediated transport | 3.70E-07 | positive regulation of cytosolic calcium ion-concentration | 4.99E-03 |
| regulation of catalytic activity | 7.49E-07 |  |  |
| biological regulation | 8.45E-07 |  |  |
| response to stimulus | 1.19E-06 |  |  |
| regulation of biological process | 1.53E-06 |  |  |

No differentially expressed genes found in this study overlapped with ISGs upregulated during acute infection identified by Bosinger and colleagues in SMs (2009) or Jacquelin and colleagues in AGMs (2009), both cercopithecine primates. Similarly, no differentially expressed genes found in this study overlapped with the immunosuppressive genes those in the Human Immunosuppressive Genes database, we identified one immunosuppressive gene, *CD101* (ENSG00000134256), that was significantly upregulated in infected individuals (fold change = 0.96; *Q* value = 0.14).

### Whole blood deconvolution – CD4+ T cell abundance

We determined the relative abundance of three CD4+ T cell subsets using computational deconvolution as implemented in CIBERSORT. The proportion of total CD4+ T cells, as a fraction of total immune cells, was 0.15, *SD* = 0.04. The proportions of three CD4+ T cell subsets as a fraction of total immune cells were significantly deconvoluted (*p* < 0.005), and abundance estimates were as follows: naïve CD4+ T cells (0.02, *SD* = 0.03), memory resting CD4+ T cells (0.02, *SD* = 0.03), and memory activated CD4+ T cells (0.10, *SD* = 0.02; Table 2). There were no significant differences in the abundance of naïve, memory resting, memory activated, or total CD4+ T cells between infected and uninfected individuals (*t*(27) = 0.29, *p* = 0.77; *t*(27) = −1.4, *p* = 0.17; *t*(27) = −0.27, *p* = 0.78; *t*(27) = −1.1, *p* = 0.29, respectively; Figure 6; full CIBERSORT results in Supplemental Table S8). We also did not detect any significant differences in CD4+ T cell subsets between males and females (*t*(27) = −0.49, *p* = 0.63; Table 2).

**Table 2.** Relative CD4+ T cell subset and total CD4+ T cell proportions (95% CI in parentheses) in infected and uninfected individuals, and males and females. P values denote no significant differences in cell proportions between infected and uninfected individuals or between males and females.

|  | Naïve CD4+ T cells | Memory resting CD4+ T cells | Memory activated CD4+ T Cells | Total CD4+ T cells |
| --- | --- | --- | --- | --- |
| SIVkrc Infected | 0.025 (0.002-0.048) | 0.016 (0.002-0.031) | 0.1 (0.088-0.112) | 0.14 (0.118-0.167) |
| SIVkrc Uninfected | 0.021 (0.007-0.036) | 0.036 (0.015-0.056) | 0.1 (0.088-0.117) | 0.16 (0.138-0.183) |
| *p* value | 0.78 | 0.17 | 0.79 | 0.30 |
| Males | 0.028 (0.012-0.045) | 0.023 (0.006-0.039) | 0.104 (0.092-0.116) | 0.156 (0.134-0.178) |
| Females | 0.009 (0.002-0.022) | 0.041 (0.019-0.062) | 0.095 (0.081-0.109) | 0.146 (0.125-0.167) |
| *p* value | 0.20 | 0.25 | 0.41 | 0.63 |
| Total | 0.023 (0.011-0.036) | 0.028 (0.015-0.041) | 0.101 (0.092-0.111) | 0.152 (0.0136-0.169) |

**Figure 6.**
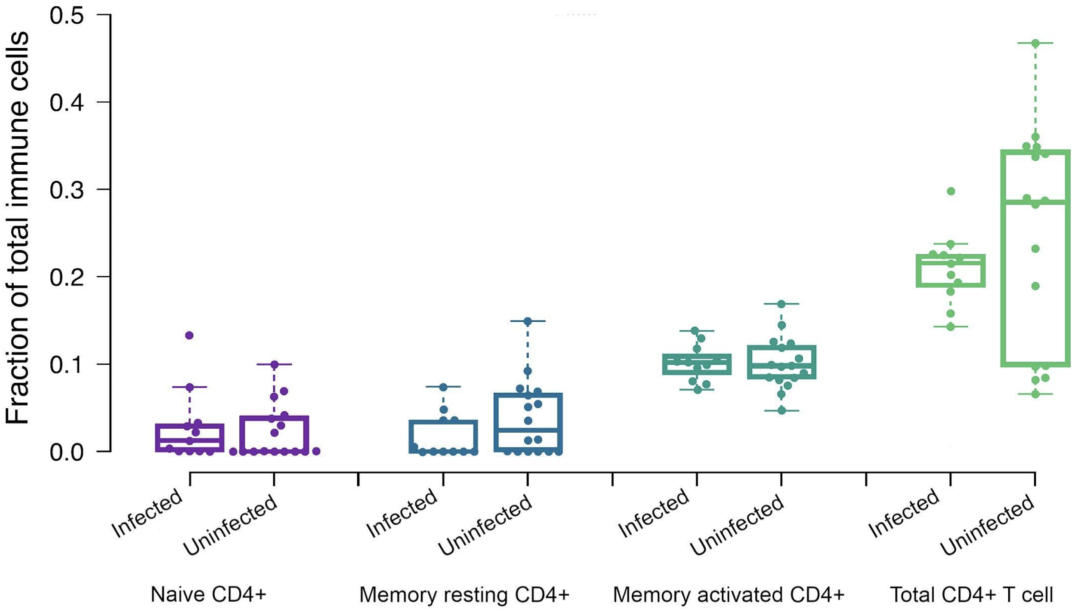
Estimated relative proportions of CD4+ T cell subsets and total CD4+ T cells from gene expression deconvolution showing no significant difference between infected and uninfected individuals.

## DISCUSSION

Nonpathogenic SIV infection in natural hosts is not adequately defined because of the narrow set of taxa in which it has been investigated. Additionally, variation has been observed in outcomes related to infection, specifically the loss of CD4+ T cells in chimpanzees and MNDs (Keele et al., 2009; Greenwood et al. 2014), and decreased birth rate, increased infant mortality risk, and reduced longevity with findings consistent with end-stage AIDS in chimpanzees (Keele et al. 2009). Due to the small range of species in which nonpathogenic SIV infection has been studied at the molecular level, the degree to which patterns of host gene expression related to SIV infection are conserved across natural hosts is unknown. Our study helps address this gap by providing novel insights into the relationship between SIV infection and genome-wide host gene expression patterns for a primate species outside of the Cercopithecinae using high-resolution RNA-seq. The main findings from our study are that 1) genome-wide gene expression patterns in SIVkrc-infected versus uninfected individuals support the hypothesis that SIVkrc has minimal effects on host immunoregulatory gene expression in URC, as we found no evidence for chronic immune gene activation or upregulated ISG expression in infected individuals, and 2) that *CD101*, an immunosuppressive gene whose downregulation is related to HIV progression, was upregulated in SIVkrc-infected URC. The upregulation of an immunosuppressive gene parallels patterns observed in AGMs and SM, although each of the three host taxa shows upregulation of different immunosuppressive genes. This finding suggests that species-specific mechanisms may underlie tolerance to SIV infection in different African primate taxa. As *CD101* has not been identified in previous studies (Bosinger et al. 2009; Jacquelin et al. 2009; Lederer et al. 2009), this finding expands our knowledge of natural host responses to SIV and highlights a potentially novel pathway to nonpathogenic SIV infection.

### Staging of SIVkrc infection in the wild URC: viral load and serology

Ma and colleagues (2013) defined AGMs as being in the acute phase of infection if they simultaneously had a high viral load and were seronegative, and in the chronic phase if they had high viral loads and were seropositive. The reasoning behind this approach is that high viral replication during acute infection generally precedes seroconversion (Fiebig et al., 2003), which can result in high viral load and seronegative results. Following this approach, we found that all of our SIVkrc infected individuals were seropositive by HIV-2 western blot testing. This finding and our finding of no evidence for upregulated ISG expression in infected individuals, strongly suggest that all infected individuals in our study were likely in the chronic phase of infection.

### Shared signatures of gene expression associated with nonpathogenic SIV infection

We hypothesized that if SIVkrc infection is nonpathogenic in URC, then URC might share similar patterns of attenuated ISG expression in the chronic phase with AGMs and SMs. More specifically, we predicted that ISG expression would not be significantly upregulated in chronically infected URC compared to uninfected URC, as AGMs and SMs attenuate ISG expression in the chronic phase (Bosinger et al. 2009; Jacquelin et al. 2009; Lederer et al. 2009). We found evidence for attenuated ISG expression, as no ISGs were significantly upregulated in SIVkrc-infected URC compared to uninfected individuals. This result was also consistent with patterns of cytokine expression (a biomarker of immune activation) in wild AGMs, where SIV-infected individuals did not differ from uninfected individuals (Ma et al. 2013).

The results of our clustering analyses (both hierarchical clustering and PCA) also support the conclusion that SIVkrc infection is nonpathogenic in URC, because infected and uninfected individuals did not form separate clusters. Because *lack* of a chronic immune response characterizes nonpathogenic infection, infection status may not be a meaningful biological distinction at the level of blood transcriptome gene expression patterns. As would be expected in blood transcriptome profiles, we observed similar signatures of immune activity (enriched immune-related GO pathways) in both infected and uninfected individuals based on K-means clustering of genes, but found no evidence of elevated immune activity in infected individuals.

### Evidence for the upregulation of immunosuppression genes in chronic phase SIV infection in natural hosts

Bosinger and colleagues (2009) and Jacquelin and colleagues (2009) found a small number of immunosuppressive genes in SMs and AGMs, respectively, that were upregulated upon infection and remained significantly upregulated through the end of each study (six months, and two years into the chronic phase of infection, respectively). The upregulated immunosuppressive genes did not overlap between the two studies, suggesting that hosts may vary in pathways activated for immunosuppression. We hypothesized that if SIVkrc infection is nonpathogenic, URC might also show upregulation of immunosuppressive genes. We did identify an important immunosuppressive gene, *CD101* (ENSG00000134256), that was significantly upregulated in infected individuals (fold change = 0.96; *Q* value = 0.14).

This finding is notable because *CD101* has not previously been identified as an upregulated immunosuppressive gene in either AGMs or SMs even though *CD101* was included in the microarrays for the previous studies, and is highly conserved across Old World monkeys. The function of *CD101* suggests it could play a role in actively down-regulating the acute immune response to SIV infection in natural hosts. Protein-coding variants in *CD101* are associated with increased susceptibility to HIV-1 (Mackelprang et al., 2017) suggesting that it plays a role in HIV infection. Additionally, reduced expression of *CD101* on mucosal CD8+ T cells in the human intestinal and pulmonary mucosa is associated with increased inflammation (Brimnes et al., 2005), demonstrating its role in immunosuppression. Perhaps most importantly, *CD101* expression is strongly associated with the suppression of T regulatory cell proliferation, a hallmark of chronic immune activation that is directly associated with progression to AIDS in HIV patients (Deeks et al., 2004; Giorgi et al., 2002; Liu et al., 1997). These previous studies, together with our finding of *CD101* upregulation in the chronic phase of SIVkrc infection, suggest that *CD101* may be an important component of nonpathogenic SIV infection in URC.

### No evidence for decrease in CD4+ T cell abundance in SIVkrc infection

In contrast to AGMs and SMs, MNDs and chimpanzees show a significant loss of memory CD4+ T cells as a consequence of SIV infection (Keele et al 2009; Greenwood et al. 2014). Because substantial variation exists between natural hosts in the effect of SIV infection on CD4+ T cell homeostasis, we were interested in whether URC would be more similar to AGMs and SMs, or MNDs and chimpanzees. We found no evidence for a decrease in abundance of any CD4+ T cell subset (naive, memory resting, or memory activated CD4+ T cell) or in total CD4+ T cell abundance in infected individuals based on deconvolution results. Given that SIVkrc and SIVmnd-1 are closely related lineages, and both SIVagm and SIVsm are distantly related, a result where URC and MNDs both showed a loss of memory CD4+ T cells in infected individuals would be consistent with a mechanism of CD4+ T cell loss driven by viral factors. Instead, our results present a more complicated picture with both host and viral factors potentially influencing the loss of activated and bystander CD4+ T cells (Doitsh & Greene, 2016).

### Potential study limitations

Our study does not perform a longitudinal assessment of the effect of SIVkrc infection on gene expression beginning in the acute phase. Because our study was conducted on a natural population, and our samples were collected cross-sectionally, we are not able to time sample collection to intentionally capture the acute response to infection. An interesting shared feature of AGMs and SMs is the robust and transient upregulation of ISGs in the acute phase of infection. While our study shows that SIVkrc-infected URC in the chronic phase do not have upregulated ISGs, it remains unknown whether the longitudinal pattern of gene expression, beginning with acute infection, is conserved in cercopithecines or more broadly conserved across African Old World monkeys. Additionally, although it appears that natural hosts differ from humans in their ability to attenuate a chronic immune response, it may be possible that older URC individuals who have been infected for a long period of time experience some level of immunodeficiency late in life, though our data are not able to address that possibility. Although the type of repeated sampling necessary to compare to experimental infection studies will not likely be possible in natural populations, the study of host responses to SIV in natural populations nonetheless adds considerably to our understanding of variation in nonpathogenic SIV infection.

Second, we focused specifically on the effects of SIVkrc infection on host gene expression. Although we control for some of the major sources of biological and technical variation in gene expression, there are many factors, both intrinsic and extrinsic (e.g. circadian patterns, age), that influence gene regulation in natural populations. Additionally, URC are host to a myriad of infectious agents including bacteria, viruses, and eukaryotic parasites (Bailey et al., 2014, 2016a, 2016b; Ghai et al., 2014; Goldberg et al., 2008; Goldberg et al., 2009; Lauck et al., 2011; Lauck et al., 2013; McCord et al., 2014; Salyer et al., 2012; Sibley et al. 2014; Simons et al., 2017; Thurber et al., 2013), all of which likely influence patterns of host gene expression. Although it is impossible to account for all of the possible factors contributing to gene expression patterns in a natural population, our study provides valuable information on the role of SIV infection in host gene expression patterns, and will inform future studies aimed at understanding co-infection dynamics.

## CONCLUSIONS

We found evidence to support the hypothesis that SIVkrc infection does not strongly influence patterns of immunological gene expression in URC based on a shared pattern of attenuated ISG expression during chronic infection with AGMs and SMs.

We found no transcriptomic evidence that individuals were immunocompromised based on the observations that 1) CD4+ T cell abundance estimates did not differ between infected and uninfected individuals (despite infected individuals harboring high viral loads), 2) infected and uninfected individuals had similar enriched GO pathways in gene clusters, and 3) infected individuals did not cluster to the exclusion of uninfected individuals using multiple clustering approaches. We did find evidence for upregulation of an important immunosuppressive gene, *CD101,* which has not been previously reported in models of nonpathogenic SIV infection, despite both protein-coding and gene regulatory variation of *CD101* being implicated in HIV infection. Based on the fundamental role of *CD101* in immunosuppression, these findings suggest evolved species-specific mechanisms for the tolerance of SIV infection in natural hosts.

## MATERIALS AND METHODS

### Ethics statement

Animal use for this study followed the guidelines of the Weatherall Report (Weatherall 2006) on the use of NHPs in research. Our study was approved by the Uganda National Council for Science and Technology, Uganda Wildlife Authority, and the University of Wisconsin-Madison and University of Oregon Animal Care and Use Committees prior to the start of the study (University of Wisconsin-Madison IACUC #A3368-01; University of Oregon IACUC #15-06A). Biomaterials were shipped from Uganda to the United States under CITES permit #002290.

### Study system and sample collection

This study is part of the Kibale EcoHealth Project, a long-term investigation of health and ecology across the human and wildlife community in and surrounding Kibale National Park, Uganda (0°13’-0°41’N, 30°19’-30°32’ E; Goldberg et al. 2012). The URC, an arboreal Old World monkey in the subfamily Colobinae, is one of the 13 primate species found in Kibale (Struhsaker 2005). The Small Camp (SC) group is a well-habituated social group that has been a focus of the Kibale EcoHealth Project since 2005 (e.g., Chapman et al. 2005, 2012; Goldberg et al. 2008, 2009; Lauck et al. 2011, 2013; Simons et al. 2016). Samples for this study were collected from 29 adult Ugandan red colobus (M = 21, F = 8) from the SC group. Sample size considerations in differential expression analysis differ from classic ecological experiments as they are a trade-off between biological replicates and sequencing depth. Our sample size (infected *n* = 12; uninfected *n* = 17) exceeded both that of similar studies (Bosinger et al. 2009; Jacquelin et al. 2009), and recommendations from a benchmarking study on sample size for detecting differentially expressed genes at all fold changes (Schurch et al. 2016). Another benchmarking study by Liu and colleagues (2014) showed diminishing returns in the power to detect differentially expressed genes beyond a sequencing depth of 10M reads per replicate. Our study also exceeded that recommendation with a minimum of 19.6M reads, and average of 23.6M per replicate (see sequencing methods below).

Animals were immobilized in the field and blood plasma was collected as previously described (Lauck et al. 2011). In addition, whole blood was collected using a modified PreAnalytiX PAXgene Blood RNA System protocol (Qiagen, Hilden, Germany). For each animal, approximately 2.5 ml of blood was collected into a PAXgene Blood RNA tube, inverted 10 times and stored at room temperature for 2-4 hours. The blood and PAXgene buffer mixture was then aliquoted into 1 mL cryovials and stored at −20°C overnight before being stored in liquid nitrogen. Samples were transported to the United States in an IATA-approved liquid nitrogen dry shipper and then transferred to −80°C for storage until further processing.

The use of whole blood was motivated by several factors: 1) it allowed comparison to the only other studies of host gene regulatory responses to nonpathogenic SIV infection, as they also used whole blood (Bosinger et al. 2009; Jacquelin et al. 2009; Lederer et al. 2009), 2) whole blood is superior to PBMCs for transcriptomic analysis when storage is required as was the case for our study (Debey-Pascher et al. 2011), 3) global transcriptomic profiles from whole blood account for latent SIV reservoirs, and may therefore be better for understanding immune responses to SIV infection (Kandathil et al. 2016), and 4) ISG gene expression profiles of SIV-infected SMs are largely concordant between whole blood and the lymph node (an important site of immune activity in response to SIV/HIV infection), enabling accurate inference of the broad transcriptomic responses to infection conserved between lymph nodes and whole blood (Bosinger et al. 2009).

### Viral prevalence and load

We assessed SIVkrc infection prevalence and viral loads using both serology and RT-qPCR. SIVkrc prevalence was assessed from serum for each individual with an HIV-2 western blot assay (MP Diagnostics HIV-2 Western Blot version 1.2; MP Biomedicals, Singapore) capable of detecting divergent SIVs (Lauck et al., 2013).

Viral load was determined by RT-qPCR detection of both SIVkrc envelope (*env*) and long terminal repeat (LTR) sequences. Viral RNA was extracted from plasma using the QIAamp Viral RNA minikit (Qiagen, Valencia, CA). For the *env* qPCR assay, the primers SIVkrc *env* F2 5’ GCA AGG TAT GAG ATC CCY AGG CA 3’ and SIVkrc *env* R4, 5’ GYA AAG CCT GGG ACT GGA TCG T 3’ and probe SIVkrc PR2 5’ FAM-AGA TAC TAA GCC CAT TGC AGT TCC “T”GC AAA GCT CAG-SpC6 3’, where “T” is labeled with BHQ1, were used to detect a 127-bp sequence. For the LTR qPCR assay, the primers SIVkrc 5LTR F 5’ GAG GTC TGT GAA GAG TGC CG 3’ and SIVkrc 5LTR R 5’ CCT TGG GGA ACT AMA CCG TC 3’ and probe SIVkrc LTR PR 5’ FAM-CTG GCA CCT CCC TG “T” ACC AGC CGC CAG TCA-SpC6 3’, where “T” is labeled with BHQ1, were used to detect a 94-bp sequence. One µl of RNA extract equivalent to 50 µl plasma was tested using the AgPath-ID One Step RT-PCR kit (Applied Biosystems, Foster City, CA) on a BioRad CFX96 iCycler (Hercules, CA) with a reverse transcriptase step at 45°C for 15 min, denaturation at 95°C for 10 min, followed by 55 cycles of 95°C and 62°C for 15 sec each. Both *env* and LTR PCR products generated from one WB-positive URC (RC51) were cloned into the pCR 2.1 TOPO vector (Thermo Fisher, Waltham, MA) and RNA standards for the qPCR assays were synthesized in vitro using the RiboMAX large scale RNA production system-T7 (Promega, Madison, WI) following the manufacturer’s instructions. Both assays could detect SIVkrc RNA in a linear range from 10^1^ to 10^7^copies/reaction and could reliably detect five viral copies/reaction. All no-template and negative plasma controls always tested non-reactive.

### RNA Extraction, Library Preparation, and Sequencing

Blood samples were thawed at room temperature for two hours, and total RNA extraction followed the PreAnalytiX PAXgene Blood RNA Kit protocol with the exception that reagents were scaled for extraction of 1 mL rather than the full PAXgene Blood tube volume. Extracts were concentrated following the manufacturers protocol for the ZYMO Research RNA Clean and Concentrator kit (Irvine, CA, USA). Alpha and beta globin mRNA was depleted from total RNA extracts following the manufacturer’s protocol for the GLOBINclear Kit (Waltham, MA, USA). Prior to library preparation, integrity of globin-depleted total RNA was assessed on an AATI Fragment Analyzer (Ankeny, IA, USA), and all 29 samples (RIN mean: 8.1, range: 6.6-9.2) were used for downstream analyses. Sequencing libraries were prepared from globin-depleted total RNA using the KAPA Biosystems Stranded mRNA-seq Kit (Wilmington, MA, USA) and indexed with Illumina adapters for multiplexing all 29 individuals on a single lane. Libraries were sequenced on three replicate Illumina NextSeq 500 high output runs with single-end 150 base pair reads at the Genomics & Cell Characterization Core Facility at the University of Oregon.

### Mapping and quantification

Demultiplexing, adapter trimming, and initial quality filtering were performed with default parameters within the Illumina NextSeq base calling software. Quality was assessed in FastQC v0.11.3 and reads for each individual were concatenated from three NextSeq runs. Additional quality filtering was performed with FASTX-toolkit, including trimming the first 15 base pairs from all reads, and removal of reads with quality score < Q20, and reads < 25 base pair length. Quality filtered RNA-seq reads were mapped to the Ugandan red colobus genome (assembly accession: GCF_002776525.1) using STAR v.2.5 in two-pass mode with -- outFilterMismatchNoverLmax 0.05. The number of reads mapping to each feature in the annotation was quantified using HTSeq v.0.9.1 (Anders et al. 2015) in intersection-nonempty mode with the –-stranded reverse option.

### Data visualization

We used multiple visualization methods to assess the degree to which SIVkrc infection separated our study subjects into distinct clusters. The feature counts matrix was pre-processed by removing low count genes for visualization. Genes with less than one count per million, calculated with CPM function in edgeR (Robinson et al. 2010), in at least three samples were removed. Missing values were treated as zeroes. The resulting dataset was then normalized using the variance stabilizing transformation (VST) as implemented in DESeq2 v.1.16.1 (Anders & Huber, 2010; Love et al., 2014) for visualization with hierarchical clustering, principal component analysis (PCA), and k-means clustering.

#### Hierarchical clustering

Hierarchical clustering was performed using the 1,000 most variable genes (based on standard deviation ranks of all genes) with sample clustering based on the distance matrix (1-*r*, where *r* is Pearson’s correlation coefficient). Hierarchical clustering was visualized as a dendrogram based on sample distances.

#### Principal component analysis (PCA)

To identify whether infected and uninfected samples clustered together we visualized our data in two-dimensional space using PCA. PCA graph was plotted using VST transformed data in R v.3.3.3 (R Core Team 2014). PCA loadings were attributed to genes (similar to expression values) and pathway analysis was conducted on the first four principal components using *pgsea* (Furge & Dykema, 2006) in R v.3.3.3. The false discovery rates (FDRs; (Benjamini & Hochberg, 1995) of significant pathways are reported for each principal component.

#### K-means clustering

K-means clustering was used to cluster groups of genes based on their expression patterns in the blood transcriptome of infected and uninfected individuals. Genes were first ranked by standard deviation, and the top 2,000 most variable genes were used for clustering. The optimal number of clusters was determined based on the elbow method, wherein the addition of additional clusters does not substantially reduce the within groups sum of squares. The number of clusters was confirmed with t-SNE mapping (Supplemental Figure S1). Enriched GO pathways were then identified for genes in each cluster using GSEA (Subramanian et al., 2005). For each cluster with significantly enriched pathways, a promoter analysis was conducted using a 300-bp region upstream of the transcription start site. For genes in significantly enriched pathways, the binding affinity of promoters to transcription factor binding sites was estimated using CIS-BP (Weirauch et al., 2014). Rather than using an arbitrary cutoff for a binary outcome of binding vs. non-binding, the best binding score for each transcription factor to every promoter sequence was used to test (Student’s t test) for transcription factor binding motifs enriched in clusters. P values of enriched transcription factor binding motifs were corrected for multiple testing and FDR is reported.

### Weighted gene co-expression network analysis

We identified co-expression networks and modules in SIVkrc-infected and uninfected individuals using weighted gene co-expression network analysis (WGCNA; Langfelder & Horvath, 2008). Co-expression networks were identified from the top 3,000 most variable genes. To reduce noise in the correlation matrix we used a soft threshold based on the smallest power at which the scale-free topology index reached 0.9, and limited the minimum module size to 20 genes. Genes in each of the modules within each network were analyzed for enriched GO biological processes using gene set enrichment analysis. Gene set enrichment analysis was performed using GSEA as implemented in iDEP.53 (Ge, 2017) with a minimum and maximum geneset size filter of 10 and 2,000, respectively, and default FDR threshold of 0.2. GSEA was run in pre-ranked mode with the GO biological process geneset background with the increased speed algorithm *fsgea* (Sergushichev, 2016).

### Differential expression analysis

We modeled the effects of SIVkrc infection on gene expression using a generalized linear model with a negative binomial distribution as implemented in DESeq2. Low count genes (e.g. those not typically expressed in blood) were not pre-filtered for differential expression analysis as DESeq2 performs independent filtering to exclude genes with low power from FDR calculations, and by default optimizes the threshold to maximize the number of differentially expressed genes given a specified FDR (Love et al. 2014), thus allowing identification of potentially important differentially expressed genes that may be expressed at low levels. We modeled the effect of SIV infection on gene expression using a pairwise design (infected N = 12 vs. uninfected N = 17) with sex and RIN score as blocking factors. P values were corrected form multiple testing and the adjusted *p* values are reported as *Q* values (Storey & Tibshirani, 2003). Because our sample size may have limited our power to detect differentially expressed genes, we followed Charruau and colleagues (2016) in setting a relatively liberal *Q* threshold of 0.2 for detecting candidate genes associated with infection. To determine possible functional enrichment in differentially expressed genes, we performed a GO term enrichment analysis in g:Profiler (Reimand et al., 2007) with minimum functional category and query size both set at 3, and with *p* values corrected using the default g:SCS threshold. We compared the differentially expressed genes to ISGs, chemokines, defensins, inhibitors, and antiviral restriction factors identified as being upregulated in acute infection and attenuated in the chronic phase in AGMs and SMs (Bosinger et al. 2009; Jacquelin et al. 2009) to assess if any of those genes were also upregulated in SIV-infected URC. We note that those two comparative studies were both performed with microarrays, and while the authors detail extensive efforts to ensure that AGM and SM hybridization to the arrays was comparable to rhesus macaque, the use of RNA-seq in our study provides an inherently unbiased view of transcriptome-wide patterns of gene expression. We also compared our differentially expressed genes to immunosuppressive genes that were upregulated in acute SIV infection and remained upregulated into the chronic phase of infection in AGMs and SMs. Finally, to identify immunosuppressive genes that might be upregulated in URC, but not AGMs or SMs, we compared our differentially expressed genes to a curated database of immunosuppressive genes in the Human Immunosuppression Gene Database (Liu et al., 2017).

### Whole blood deconvolution of immune cell subsets

The effect of SIV infection on CD4+ T cell maintenance varies across AGMs, SMs, MNDs, and chimpanzees. While AGMs and SMs maintain CD4+ T cell homeostasis, naturally infected MNDs and chimpanzees show a loss of memory CD4+ T cells (Keele et al. 2009; Greenwood et al. 2014). We obtained whole blood from a natural population in a remote field site and could therefore not perform immunohistochemistry and flow cytometry to isolate T cell subsets. To test the effect of SIVkrc infection on CD4+ T cell levels we estimated the relative abundance of immune cell subsets using the computational deconvolution method implemented in CIBERSORT (Newman et al., 2015). This is a robust method for determining the relative proportion of immune cell subsets (including T cells, memory B cells, natural killer (NK) cells, monocytes, macrophages, and neutrophils), and is superior to other deconvolution methods in accuracy, sensitivity to noise, and resolution of closely related cell types, particularly for whole blood deconvolution (Newman et al., 2015; Wang et al., 2016). CIBERSORT was run with 1,000 permutations, and accurate deconvolution of cell types for each sample was assessed with goodness of fit. Differences in relative fractions of naïve, memory resting, memory activated, and total CD4+ T cells between SIVkrc-infected and uninfected individuals were tested with a two-sample t-test.

## ACKNOWLEDGEMENTS

This research was funded by NIH grant TW009237 as part of the joint NIH–NSF Ecology of Infectious Disease program and the UK Economic and Social Research Council, NSF BCS-1540459, National Geographic Society, NSERC, and the University of Oregon. We thank the Uganda Wildlife Authority and Uganda National Council of Science and Technology for permission to conduct this research. We are grateful to Robert Basaija, Peter Tuhairwe, Clovice Kaganzi, and Dr. Dennis Twinomugisha for assistance with logistics and fieldwork. We thank Clay Small, Lucia Carbone, Suzi Fei, Dave O’Connor, and Tom Friedrich for helpful feedback on the design and analysis of the research and members of the Molecular Anthropology Group at University of Oregon for helpful feedback throughout the research process and valuable comments during manuscript preparation. Use of trade names is for identification only and does not imply endorsement by the U.S. Centers for Disease Control and Prevention (CDC). The findings and conclusions in this report are those of the authors and do not necessarily represent the views of the CDC.

David Hyeroba, our co-author, colleague, and friend, passed away while we were in the late stages of submitting this manuscript for publication. Dr. Hyeroba was an incredible and passionate veterinarian who worked internationally to care for the health of wild and domestic animals. This manuscript would not have been possible without his expertise and skill, and his passing is an irreparable loss to our community. We dedicate this research to his memory.

